# Cholesterol binds the amphipathic helix of IFITM3 and regulates antiviral activity

**DOI:** 10.1101/2022.04.21.488780

**Authors:** Kazi Rahman, Siddhartha A.K. Datta, Andrew H. Beaven, Abigail A. Jolley, Alexander J. Sodt, Alex A. Compton

## Abstract

The interferon-induced transmembrane (IFITM) proteins broadly inhibit the entry of diverse pathogenic viruses, including Influenza A virus (IAV), Zika virus, HIV-1, and SARS coronaviruses by inhibiting virus-cell membrane fusion. IFITM3 was previously shown to disrupt cholesterol trafficking, but the functional relationship between IFITM3 and cholesterol remains unclear. We previously showed that inhibition of IAV entry by IFITM3 is associated with its ability to promote cellular membrane rigidity, and these activities are functionally linked by a shared requirement for the amphipathic helix (AH) found in the intramembrane domain (IMD) of IFITM3. Furthermore, it has been shown that the AH of IFITM3 alters lipid membranes *in vitro* in a cholesterol-dependent manner. Therefore, we aimed to elucidate the relationship between IFITM3 and cholesterol in more detail. Using a fluorescence-based *in vitro* binding assay, we found that a peptide derived from the AH of IFITM3 directly interacted with the cholesterol analog, NBD-cholesterol, while other regions of the IFITM3 IMD did not, and native cholesterol competed with this interaction. In addition, recombinant full-length IFITM3 protein also exhibited NBD-cholesterol binding activity. Importantly, previously characterized mutations within the AH of IFITM3 that strongly inhibit antiviral function (F63Q and F67Q) disrupted AH structure in solution, inhibited cholesterol binding *in vitro*, and restricted bilayer insertion *in silico*. Our data suggest that direct interactions with cholesterol may contribute to the inhibition of membrane fusion pore formation by IFITM3. These findings may facilitate the design of therapeutic peptides for use in broad-spectrum antiviral therapy.

## Introduction

Interferon-induced transmembrane (IFITM) proteins belong to a family of small transmembrane proteins known to interfere with diverse membrane fusion events important for human physiology (Coomer et al., 2021; Shi et al., 2017). IFITM proteins inhibit the cellular entry step of many enveloped viruses pathogenic to humans, like Influenza A virus (IAV), Ebola virus (EBOV), Dengue virus (DENV), SARS coronaviruses, and HIV-1 (Majdoul and Compton, 2021). IFITM3 is the most characterized family member owing to its antiviral potency and its links to genetic susceptibility to infection in human populations (Zhao et al., 2018). Previous research has shown that IFITM3 inhibits the virus-cell membrane fusion process by blocking fusion pore formation, a terminal step of the entry process that enables access of enveloped virions to the host cell cytoplasm (Desai et al., 2014; Li et al., 2013). We recently demonstrated that an amphipathic helix (AH) in the amino terminus of IFITM3 (Chesarino et al., 2017) confers IFITM3 with the ability to alter the biomechanical properties of membranes in living cells (Rahman et al., 2020). Specifically, IFITM3 decreases membrane fluidity (increases membrane rigidity) in living cells in an AH-dependent manner (Rahman et al., 2020). This was confirmed *in vitro* by showing that a peptide corresponding to the AH of IFITM3 is sufficient to promote membrane rigidity in artificial membranes, and interestingly, the AH requires membrane cholesterol in order to alter membranes (Guo et al., 2021). Furthermore, a sterol binding antibiotic, Amphotericin B, negates the antiviral activity of IFITM3 (Lin et al., 2013) and prevents membrane stiffening by IFITM3 (Rahman et al., 2020). These reports indicate that the AH and cholesterol are important for the functions of IFITM3, but the relationship between them is poorly understood.

Establishing how IFITM3 interacts with and influences the membrane microenvironment is key to understanding how it inhibits membrane fusion pore formation. Cholesterol is a key regulator of the biomechanical properties of lipid bilayers and is known to influence the cell entry step of enveloped viruses (Chernomordik and Kozlov, 2003; Teissier and Pécheur, 2007). A link between IFITM3 and cholesterol was first raised by showing that IFITM3 disrupts the function of VAMP-associated Protein A (VAPA), a protein controlling cholesterol transport between the endoplasmic reticulum and late endosomes/multivesicular bodies (Amini-Bavil-Olyaee et al., 2013). As a result, IFITM3 triggers cholesterol accumulation within late endosomes. However, the relevance of this phenotype to the antiviral mechanism of IFITM3 remains unclear. Some studies have shown that inhibition of the cholesterol transporter NPC1, which results in intraendosomal accumulation of cholesterol, blocks infection by IAV, EBOV and DENV at the entry stage (Carette et al., 2011; Kühnl et al., 2018; Poh et al., 2012). However, other studies have reported that cholesterol redistribution to late endosomes is not sufficient to phenocopy the block to infection mediated by IFITM3 (Desai et al., 2014; Lin et al., 2013; Wrensch et al., 2014). Therefore, it is possible that both IFITM3 and cholesterol must be present in the same membranes for virus entry inhibition to occur.

In this report, we reconcile previously conflicting pieces of evidence by showing that IFITM3 directly interacts with cholesterol. This interaction is dictated primarily by the AH, but a downstream cholesterol recognition motif (CARC) also contributes to cholesterol binding potential. We show that previously described loss-of-function mutations in the AH of IFITM3 disrupt helix formation and result in loss of cholesterol binding. These findings allow for an updated model of antiviral function for IFITM3, one in which the interaction of IFITM3 with its lipid environment alters the biomechanical properties of membranes to disfavor fusion pore formation at membranes serving as entry portals for virus infection.

## Results

### IFITM3 interacts with NBD-cholesterol and binding maps to the AH

We generated peptides corresponding to regions of IFITM3, including the intramembrane domain (IMD) and the cytoplasmic intracellular loop (CIL), and tested them for cholesterol binding potential by measuring fluorescence spectroscopy of NBD-cholesterol (Wustner, 2007) (**Figure 1A**). Following excitation at 470 nm, this sterol analog emits fluorescence when bound by protein or peptide, but it does not in the unbound state. We mixed increasing concentrations of peptide with a fixed amount of NBD-cholesterol (500 nM) in NP-40-containing buffer, which is inferior to its critical micelle concentration (700 nM) (Avdulov et al., 1997). Therefore, under these conditions, NBD-cholesterol is not predicted to form micelles. As a result, NBD fluorescence most likely indicates a direct interaction between NBD-cholesterol and peptide in solution.

**Figure 1:**
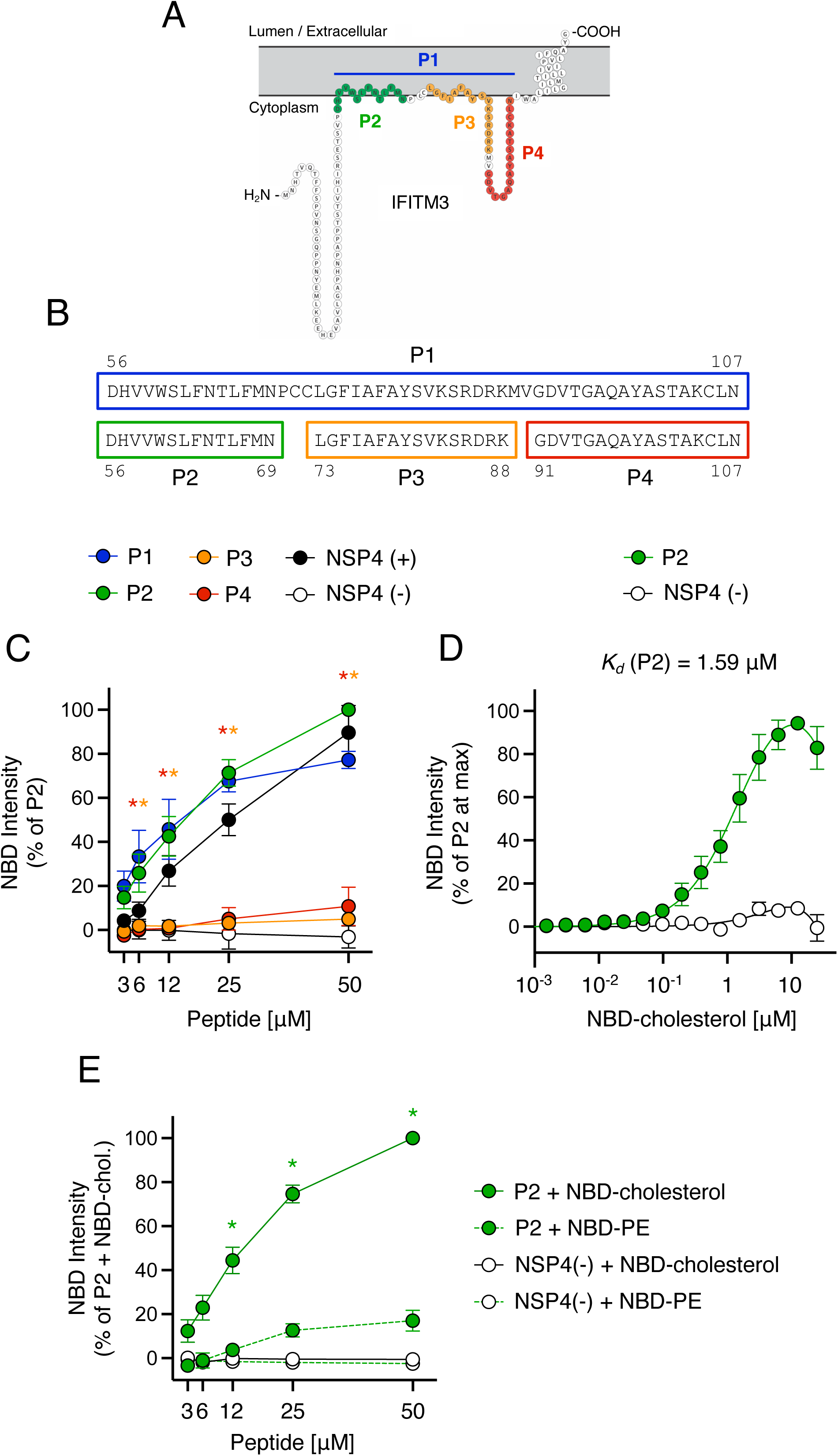
Peptides containing the AH of IFITM3 interact with NBD-cholesterol. (A) Schematic representation of the membrane topology of IFITM3 made with Protter (Omasits et al., 2014). Residues covered by indicated peptides are color-coded. (B) Detailed view of peptide sequences. Numbers correspond to amino acid residues in the context of full-length IFITM3. (C) NBD-cholesterol (500 nM) fluorescence intensities measured by spectroscopy (excitation: 470 nm; emission: 540 nm) following 16-hour incubation with increasing concentrations (3.125 to 50 μM) of peptides derived from IFITM3 or control peptides. Results represent the mean of three independent experiments and are normalized to 50 μM P2 peptide + NBD-cholesterol (set to 100%). (D) Fluorescence intensities are shown following one hour incubation of increasing concentrations of NBD-cholesterol (0.00038 to 25 μM) with 5 μM P2 or control peptide. Results represent the mean of four independent experiments and are normalized to the concentration of NBD-cholesterol that achieved maximum fluorescence in each individual experiment (set to 100%). Mean values were fitted to a nonlinear regression curve to derive the dissociation constant (*K*_*d*_). (E) Fluorescence intensities are shown for NBD-cholesterol or NBD-PE following incubation with increasing concentrations of P2 peptide or control peptide. Results represent the mean of three independent experiments and are normalized to 50 μM P2 peptide + NBD-cholesterol (set to 100%). Error bars indicate standard error. Asterisks indicate statistically significant difference (p<0.05) between P2 and another peptide (condition indicated by asterisk color; control peptides excluded), as determined by one-way ANOVA. Chol.; cholesterol. See also Table 1.

As positive and negative controls for NBD-cholesterol binding, we used peptides derived from rotavirus NSP4 that were previously shown to possess or lack cholesterol binding potential (referred to as NSP4 (+) or NSP4 (-), respectively) (**Table 1**) (Schroeder et al., 2012). Relative to these controls, we found that a peptide spanning the IMD and CIL domains of IFITM3 (amino acids 56-107, referred to as P1) enhanced NBD-cholesterol fluorescence intensity in a dose-dependent manner (**Figure 1B-C**). To map the region capable of NBD-cholesterol binding, we generated smaller peptides covering the IMD (referred to as P2 and P3) or the CIL (referred to as P4) (**Table 1 and Figure 1B**). Compared to P1, NBD-cholesterol fluorescence intensity was enhanced by P2 to a similar extent while P3 or P4 had no significant effect (**Figure 1C**). These results demonstrate that the region conferring cholesterol binding potential is found within a portion of the IMD of IFITM3 corresponding to amino acids 56-69. Notably, this region encompasses the AH of IFITM3 (defined as amino acids 59-68 (Chesarino et al., 2017)). Binding between P2 and NBD-cholesterol was apparent following one hour of incubation and the measurement was robust up to at least 16 hours (**Supplemental Figure 1**).

**Table 1:**
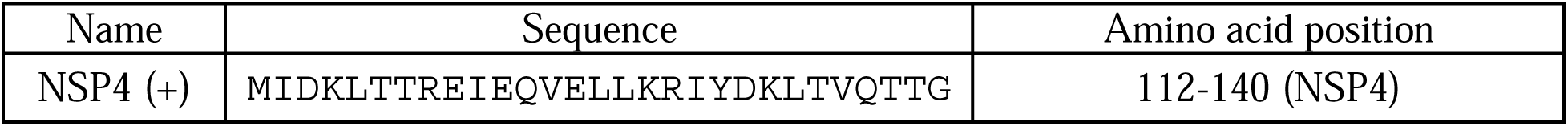
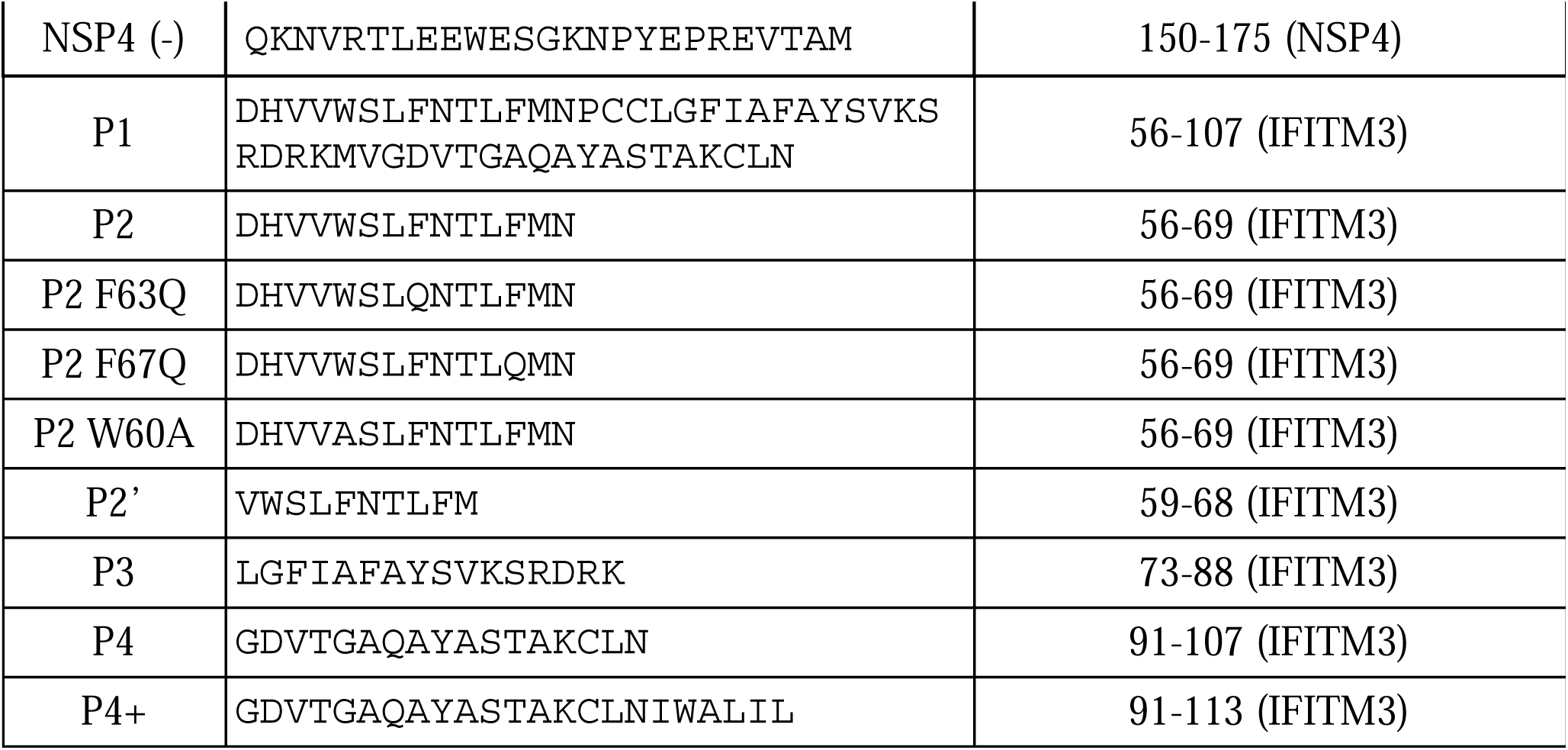
Peptides used in this study.

To measure the binding affinity between P2 and NBD-cholesterol and to exclude the possibility that increasing concentrations of peptide caused NBD-cholesterol fluorescence through a non-specific, aggregation-based mechanism, we incubated increasing concentrations of NBD-cholesterol with a fixed concentration of P2. This approach allowed us to achieve saturation of NBD-cholesterol fluorescence and to derive an apparent dissociation constant (*K*_*d*_) of 1.59 µM (**Figure 1D**). In contrast to NBD-cholesterol, the fluorescence of NBD-phosphatidylethanolamine (PE) was not significantly enhanced by P2 (**Figure 1E**), suggesting that lipid binding by the AH of IFITM3 is selective for cholesterol.

To confirm that full-length IFITM3 protein also displays cholesterol binding potential *in vitro*, we assessed the capacity for recombinant GST-tagged human IFITM3 to enhance NBD-cholesterol fluorescence intensity. Accordingly, GST-IFITM3 produced a dose-dependent increase in NBD-cholesterol fluorescence, while recombinant GST alone had no effect (**Figure 2A-B**).

**Figure 2:**
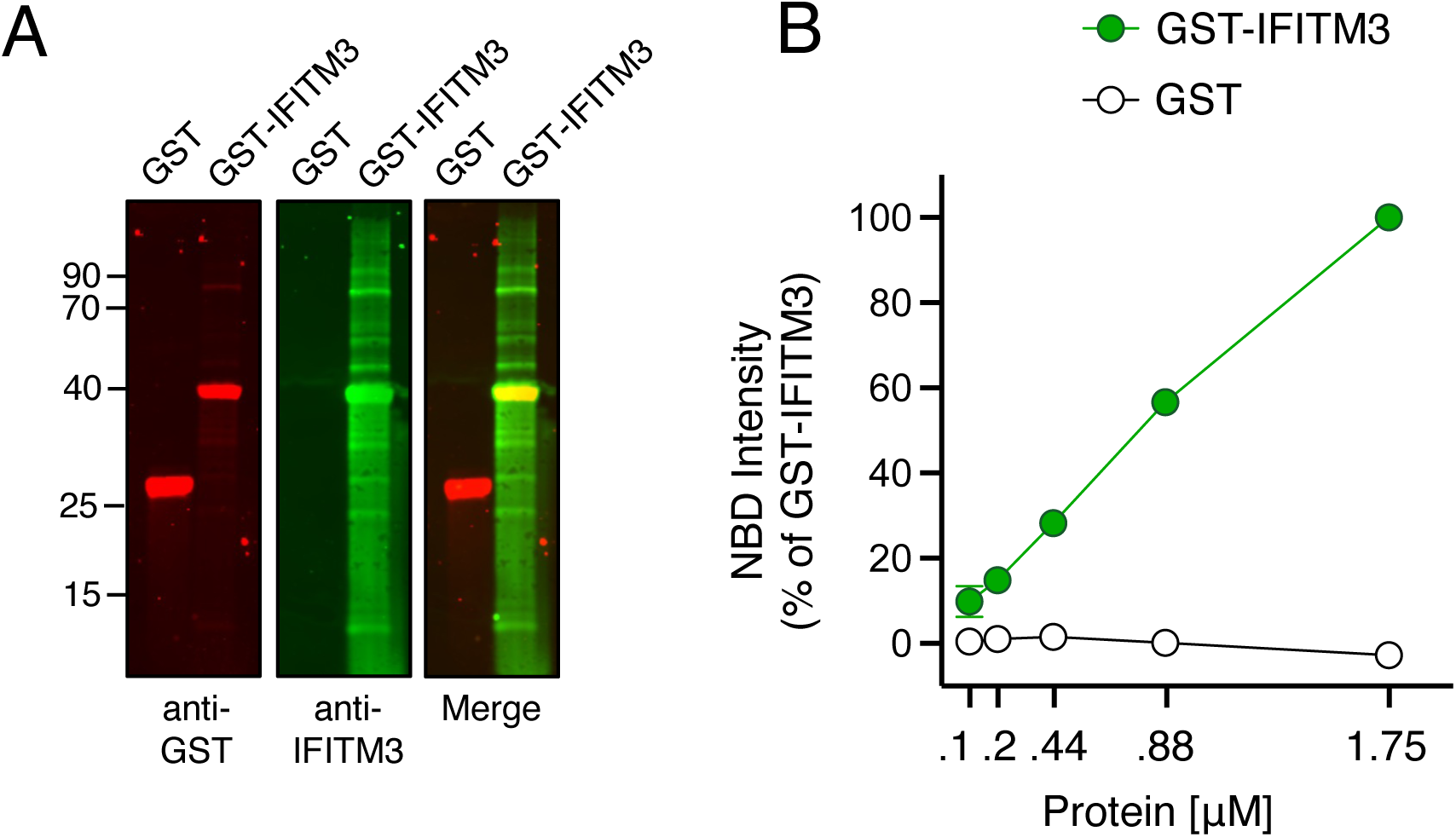
Recombinant IFITM3 protein interacts with NBD-cholesterol. (A) 2 μg of GST or GST-IFITM3 were subjected to SDS-PAGE and Western blot analysis. Immunoblotting was performed with anti-GST and anti-IFITM3 on the same nitrocellulose membrane. Numbers and tick marks indicate size (kilodaltons) and position of protein standards in ladder (ladder not shown). (B) NBD-cholesterol (500 nM) fluorescence intensities measured by spectroscopy (excitation: 470 nm; emission: 540 nm) following incubation with increasing concentrations(0.11 to 1.75 μM) of GST-IFITM3 or GST for one hour. Results represent the mean of two independent experiments and are normalized to 1.75 μM GST-IFITM3 + NBD-cholesterol (set to 100%). Error bars indicate standard error.

As a complementary approach to measuring peptide-lipid interactions, we assessed how intrinsic tryptophan fluorescence (Vivian and Callis, 2001) of P2 was affected by the presence of NBD-cholesterol or NBD-PE. P2 contains a single tryptophan at amino acid 60 (W60), and fluorescence emission was detected by spectroscopy. In contrast, mutant P2 containing W60A was not fluorogenic (**Figure 3A**). NBD-cholesterol has a minor excitation peak between 300 and 400 nm, and thus may absorb energy emitted by tryptophan as a result of Forster Resonance Energy Transfer (FRET) (Loura et al., 2001). Accordingly, we found that intrinsic tryptophan fluorescence of P2 resulted in FRET to NBD-cholesterol, while FRET to NBD-PE was minor (**Figure 3B**). This was accompanied by a decrease in tryptophan fluorescence of P2 in the presence of NBD-cholesterol, but not in the presence of NBD-PE (**Figure 3C**). These results are strongly suggestive of a direct and selective interaction between P2 and cholesterol.

**Figure 3:**
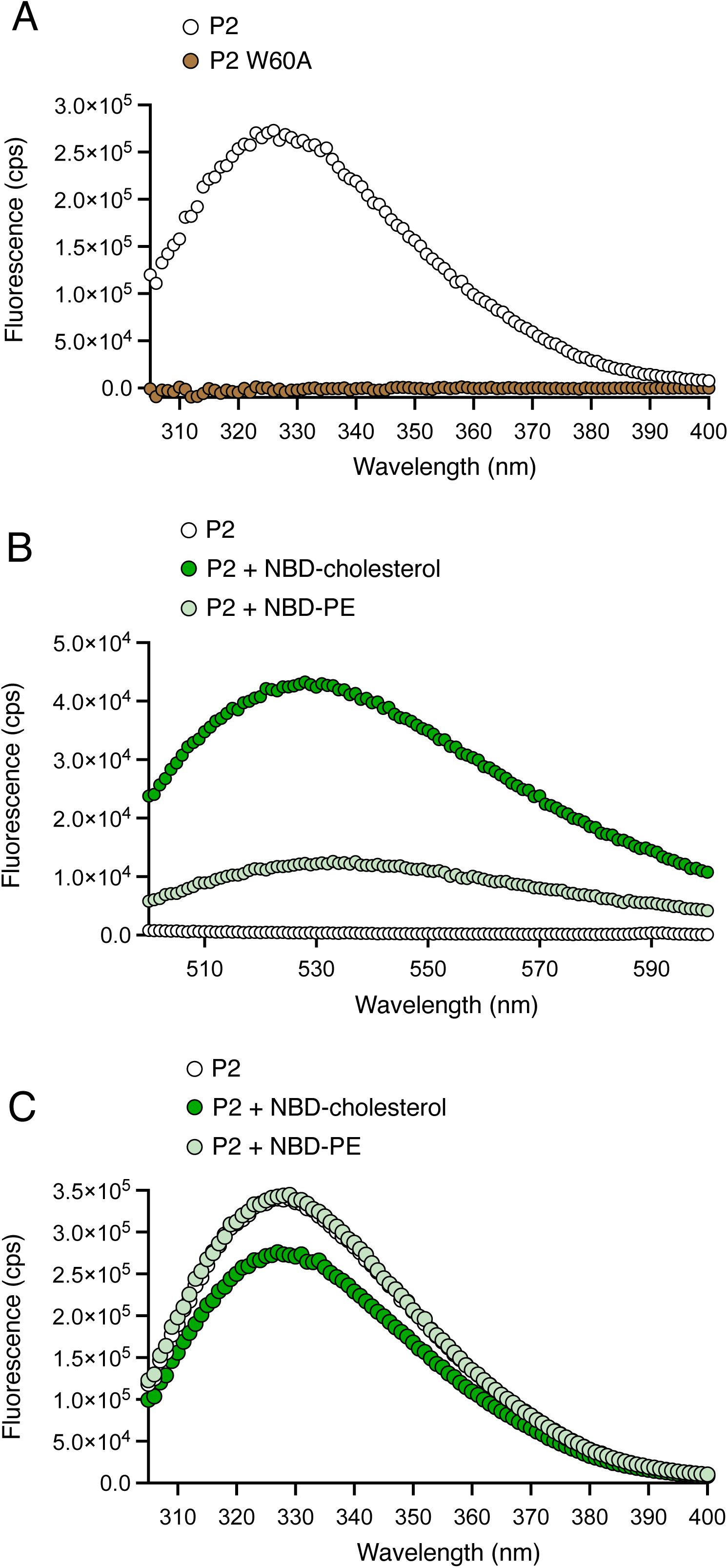
Tryptophan fluorescence of peptide containing the AH of IFITM3 transfers to NBD-cholesterol by FRET. (A) Intrinsic tryptophan fluorescence spectra for 25 μM P2 and P2 W60A peptides following excitation at 295 nm. Fluorescence emission was measured in the range of 205-300 nm. (B) 50 μM P2 in the absence or presence of NBD-cholesterol/NBD-PE was excited at 295 nm and fluorescence was measured in the range of 500-600 nm to assess FRET to NBD-cholesterol/NBD-PE. (C) 50 μM P2 in the absence or presence of NBD-cholesterol/NBD-PE was excited at 295 nm and fluorescence was measured in the range of 205-300 nm. Cps; counts per second.

### Mutations in IFITM3 that disrupt AH formation inhibit cholesterol binding and inhibit membrane insertion

To identify specific determinants for cholesterol binding within the AH of IFITM3, we introduced mutations into the P2 peptide (**Figure 4A**). F63Q and F67Q correspond to previously characterized mutations that result in near-complete loss of antiviral activity against IAV (Chesarino et al., 2017). Furthermore, F63Q was shown to abolish the capacity for an IFITM3 peptide to alter membranes *in vitro* (Guo et al., 2021). We confirmed that IFITM3 containing F63Q or F67Q exhibited a significant loss in antiviral activity against IAV compared to IFITM3 WT following transfection into cells, while IFITM3 lacking the entire AH (Δ59-68) was completely inactive (**Supplemental Figure 2A-B**). We found that, compared to P2 peptide containing the WT AH of IFITM3, introduction of F63Q or F67Q strongly reduced NBD-cholesterol binding (**Figure 4B**). To confirm that these findings are most likely the result of direct peptide-cholesterol interactions, we examined fluorescence polarization of NBD-cholesterol as a function of peptide binding. Fluorescence polarization is performed by exciting reaction mixtures with plane-polarized light and recording the degree of depolarization of emitted light. Small fluorescent molecules in aqueous solution rotate or “tumble” very quickly and assume various orientations, resulting in a high degree of depolarization of emitted light. However, upon association with a larger intermolecular complex, rotation and orientation changes are reduced, and emitted light is more polarized in nature. Consistently, P2 enhanced NBD-cholesterol fluorescence polarization in a dose-dependent manner while peptides containing F63Q or F67Q only did so modestly (**Figure 4C**). These results suggest that phenylalanines within the AH of IFITM3 are crucial determinants for cholesterol binding. Experiments performed with decreased concentrations of peptide and decreased concentration of NBD-cholesterol (50 nM) resulted in the same differential pattern of cholesterol binding among peptides, further ruling out a confounding effect of non-specific peptide-micelle interactions (**Supplemental Figure 2C**). We also produced a shorter version of the P2 peptide (referred to as P2’) that consists solely of the AH of IFITM3 (**Figure 4A**) and we observed a similar enhancement of NBD-cholesterol fluorescence polarization (**Figure 4D**). In a competition-based assay, we pre-incubated P2’ with excess unmodified cholesterol prior to adding NBD-cholesterol. We found that unmodified cholesterol competed with NBD-cholesterol for binding to P2’, resulting in a partial inhibition of NBS-cholesterol fluorescence (**Supplemental Figure 2D**). The partial competition of NBD-cholesterol binding by excess unmodified cholesterol may result from incomplete solubility of the latter under the conditions tested. Nonetheless, this finding suggests that P2’ exhibits the capacity to bind not only the cholesterol analog but native cholesterol as well. Collectively, these results further refine the cholesterol binding footprint of IFITM3 to amino acids 59-68 in the IMD (the AH itself).

**Figure 4:**
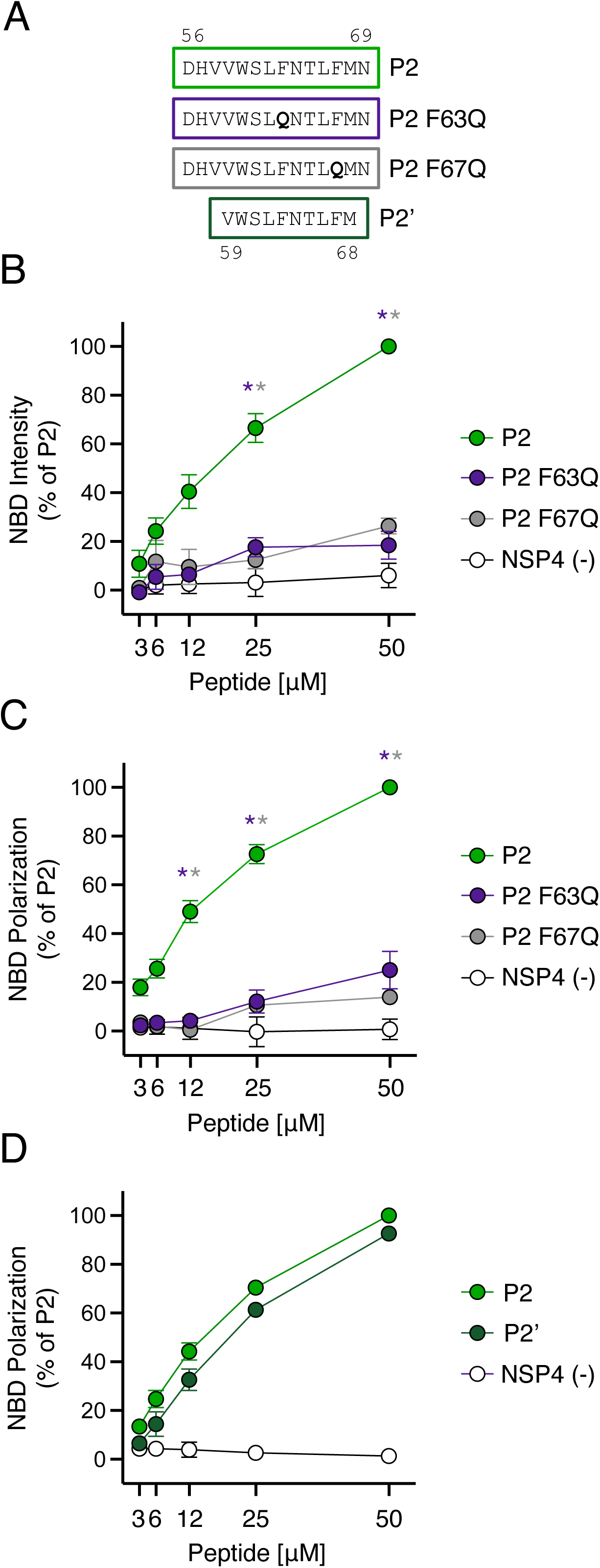
Loss-of-function mutations in IFITM3 disrupt cholesterol binding. (A) Detailed view of peptide sequences. Numbers correspond to amino acid residues in the context of full-length IFITM3. (B) NBD-cholesterol (500 nM) fluorescence intensities measured by spectroscopy (excitation: 470 nm; emission: 540 nm) following incubation with increasing concentrations (3.125 to 50 μM) of indicated peptides. Results represent the mean of three independent experiments and are normalized to 50 μM P2 peptide + NBD-cholesterol (set to 100%). (C) Fluorescence polarization of NBD-cholesterol (500 nM) measured by spectroscopy (excitation with plane-polarized light: 470 nm; emission: 540 nm) following incubation with increasing concentrations (3.125 to 50 μM) of indicated peptides. Results represent the mean of three independent experiments and are normalized to 50 μM P2 peptide + NBD-cholesterol (set to 100%). (D) Fluorescence polarization of NBD-cholesterol (500 nM) following incubation with increasing concentrations (3.125 to 50 μM) of P2, P2’, or control peptide. Results represent the mean of three independent experiments and are normalized to 50 μM P2 peptide + NBD-cholesterol (set to 100%). Error bars indicate standard error. Asterisks indicate statistically significant difference (p<0.05) between P2 and another peptide (condition indicated by asterisk color), as determined by one-way ANOVA.

To better understand how F63Q and F67Q interfere with the cholesterol binding potential of the AH, we assessed the impact of these mutations on peptide secondary structure using circular dichroism (CD). The CD spectra obtained for P2 and P2’, which possessed a similarly high capacity for NBD-cholesterol binding (**Figure 4D**), are consistent with substantial alpha-helical character (**Figure 5A**), in agreement with previous findings (Chesarino et al., 2017). Secondary structure content analysis revealed that P2 and P2’ exhibited 44% and 37% helix content, respectively (**Figure 5B**). In contrast, P2 containing F67Q presented 17% helix content and P2 containing F63Q exhibited no helical signature whatsoever (**Figure 5B**). These results suggest that the F63Q and F67Q mutations prevent proper folding of the AH, which may suggest that an intact and properly oriented helix are required for cholesterol binding.

**Figure 5:**
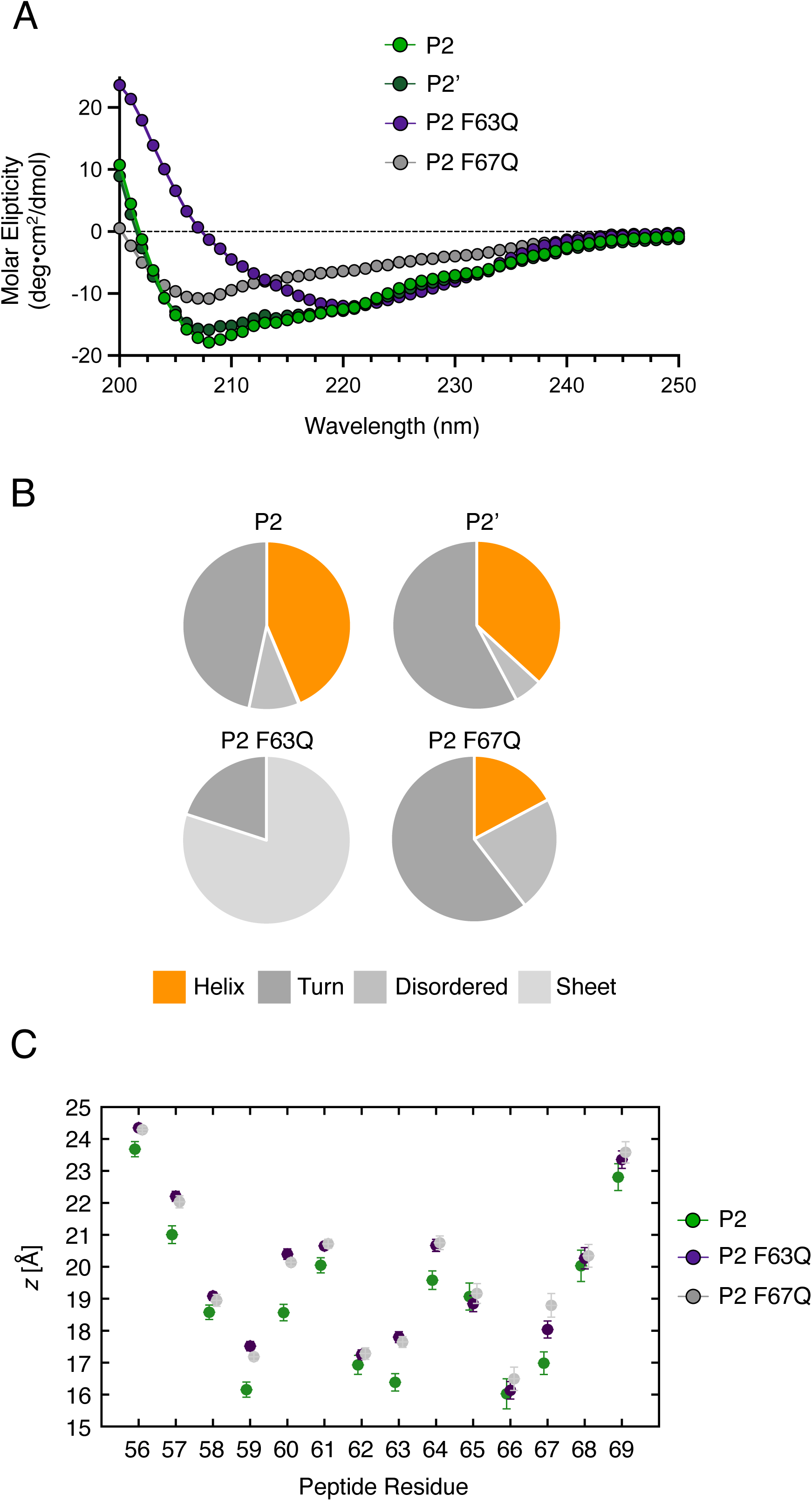
Loss-of-function mutations in IFITM3 result in loss of amphipathic helix formation and decreased membrane insertion depth. (A) Structural characterization of peptides by circular dichroism was performed in the presence of SDS. Spectra of molar ellipticity are shown for the indicated peptides across wavelengths of 200-250 nm. (B) Secondary structure content was measured using PEPFIT and presented as pie chart percentages. Averages of the top five deconvolved estimates for each peptide are presented. (C) Average alpha-carbon depth of each residue of the P2 peptide is plotted in relation to a membrane leaflet. Each leaflet is positioned such that the POPC tail termini are at *z* = 0 Å, *sn*-2 carbonyl carbon atoms are at *z =* 18 Å, and phosphorous atoms are at *z* = 22 Å. Error bars represent standard error, calculated assuming peptides in opposite leaflets are independent. Averaging the insertion depth across all peptide residues yields: WT = 19.0 ± 0.3 Å; F63Q = 19.8 ± 0.1 Å; and F67Q = 19.8 ± 0.2 Å. Å; angstroms.

Next, we used all-atom molecular dynamics simulations to model peptide insertion into a model membrane mimicking a eukaryotic lipid bilayer consisting of phosphatidylcholine (POPC) and cholesterol. Peptide depth was quantified by measuring the location of the peptide’s alpha-carbon atoms (i.e. the first carbon of each amino acid branched from peptide backbone) relative to phospholipid tail termini (the bilayer midplane, defined as *z* = 0 Å). As such, a larger *z* indicates peptide bound closer to the leaflet surface while a smaller *z* indicates peptide closer to the leaflet interior. We found that insertion of P2 WT peptide was significantly deeper than peptides containing F63Q or F67Q (**Figure 5C**). Specifically, F63Q or F67Q resulted in more shallow membrane associations at amino acid residues 56, 57, 59, 60, 61, 63, 64, and 67. Since membrane insertion depth correlates with the capacity for amphipathic helices to alter membrane order and curvature (Drin and Antonny, 2010; Sodt and Pastor, 2014), these findings provide a possible mechanistic explanation for why full-length IFITM3 protein containing F63Q or F67Q exhibits loss of function in cells and in artificial membranes (Chesarino et al., 2017; Guo et al., 2021) (**Supplemental Figure 2B**).

### A CARC motif near the TMD of IFITM3 may exhibit NBD-cholesterol binding activity

During the preparation of this manuscript, a pre-print was posted that characterized IFITM3 as a sterol-binding protein (Das et al., 2021). The authors found that endogenous IFITM3 is among the suite of host proteins that cross-links with a cholesterol analog in human cells. Furthermore, they proposed that a region of IFITM3 proximal to the transmembrane domain (TMD) encodes a putative CARC consisting of ^104^<u>K</u>CLNI<u>W</u>ALI<u>L</u>^113^ (underlined residues indicate the basic, aromatic, and aliphatic residues that define a putative cholesterol binding region, as seen in certain G-protein coupled receptors (Fantini and Barrantes, 2013)). Deletion of this region led to partial loss of cholesterol analog binding, suggesting that this region of IFITM3 protein contributes to cholesterol binding *in vivo* (Das et al., 2021). Therefore, we tested whether a peptide overlapping with ^104^<u>K</u>CLNI<u>W</u>ALI<u>L</u>^113^ conferred potential for direct cholesterol binding *in vitro*. As shown in Figure 1, P4 peptide exhibits little to no cholesterol binding activity **(Figure 1A-B)**. However, the inclusion of the putative CARC motif in P4+ led to increased cholesterol binding relative to P4 **(Figure 6A-C)**. Therefore, the ^104^<u>K</u>CLNI<u>W</u>ALI<u>L</u>^113^ motif in IFITM3 may also contribute cholesterol binding potential to IFITM3. Nonetheless, the impact of P4+ on NBD-cholesterol fluorescence is modest compared to that of P2 (**Figure 6C**).

**Figure 6:**
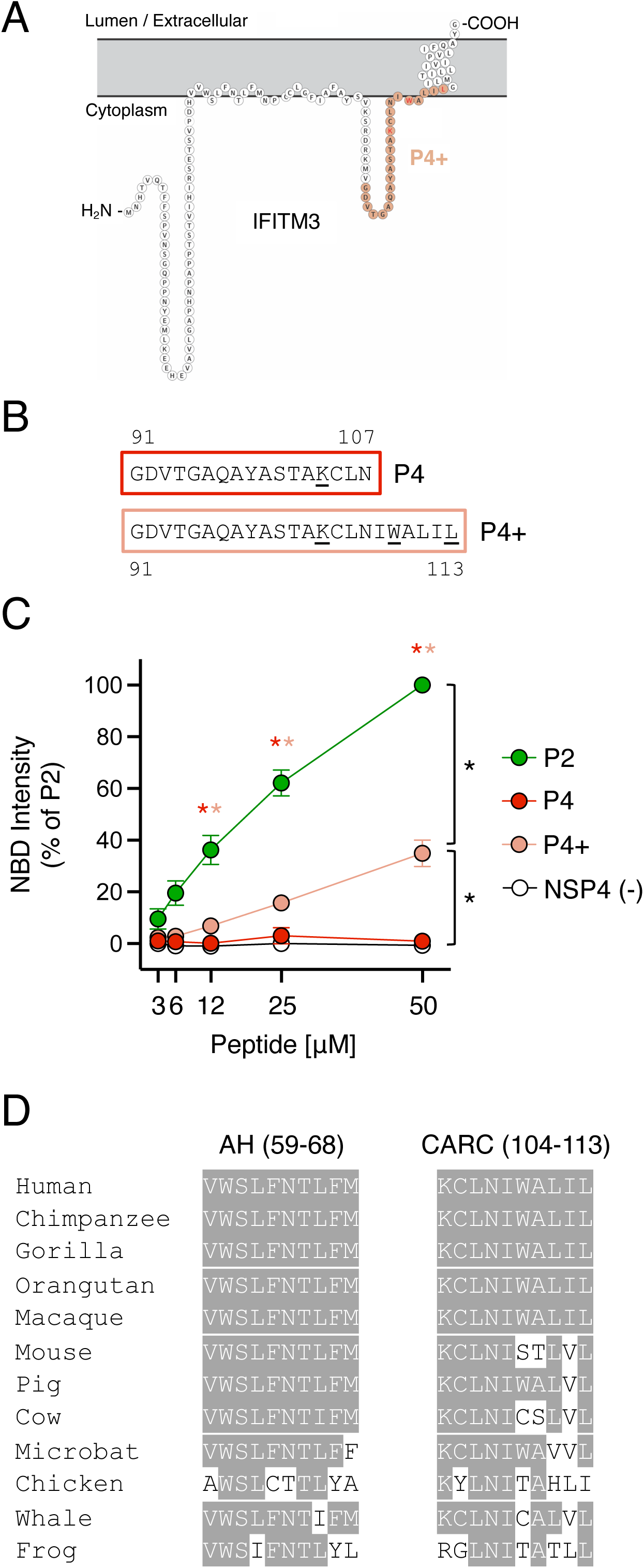
A CARC motif near the TMD of IFITM3 exhibits minor NBD-cholesterol binding activity. (A) Schematic representation of the membrane topology of IFITM3 made with Protter. Residues covered by the P4+ peptide is indicated in pink. (B) Detailed view of P4 and P4+ peptide sequences. Numbers correspond to amino acid residues in the context of full-length IFITM3. (C) NBD-cholesterol (500 nM) fluorescence intensities measured by spectroscopy (excitation: 470 nm; emission: 540 nm) following incubation with increasing concentrations (3.125 to 50 μM) of the indicated peptides. Results represent the mean of three independent experiments and are normalized to 50 μM P2 peptide + NBD-cholesterol (set to 100%). Error bars indicate standard error. Asterisks indicate statistically significant difference (p<0.05) between P2 and another peptide (condition indicated by asterisk color), as determined by one-way ANOVA. Additional pairwise comparisons also resulted in statistically significant differences between indicated pairs (black asterisks). (D) Partial amino acid alignment of IFITM3 from select vertebrate species (common names are shown). Residues conserved with human IFITM3 are highlighted in black.

Sequence analysis of *IFITM3* orthologs in diverse vertebrate species revealed that the AH is more highly conserved than the CARC (^104^<u>K</u>CLNI<u>W</u>ALI<u>L</u>^113^) region (**Figure 6D**). Notably, the latter is not conserved between human and mouse IFITM3. Since IFITM3 performs important antiviral activities in both species *in vivo* (Bailey et al., 2012; Everitt et al., 2012), this CARC motif may not be essential for antiviral function. Therefore, while our data suggest that IFITM3 contains at least two membrane-proximal regions capable of interacting with cholesterol, the cholesterol-binding activity of the AH may be the most functionally influential.

## Discussion

By showing that IFITM3 exhibits direct cholesterol binding potential via at least two membrane proximal domains (the AH and a CARC motif near the TMD), our results suggest that IFITM3 interacts with this lipid during its transit through and residency within cellular membranes. Indeed, endogenous IFITM3 has been shown to associate with a cholesterol analog in cells following cross-linking (Das et al., 2021). However, since cross-linking is not necessarily indicative of a direct interaction, our results showing the direct binding of cholesterol by IFITM3-derived peptides *in vitro* provides an important confirmation of this phenomenon but also identified the protein domain(s) responsible. Since we found that the AH is a major contributor to the direct cholesterol binding potential of IFITM3, this discovery provides a considerable leap forward in our efforts to build a molecular model for how IFITM3 inhibits fusion pore formation between viral and cellular membranes. Previous work from our laboratory and others has shown that the AH is critical for the antiviral functions of IFITM3, and that it is alone responsible for membrane alterations that block virus entry (rigidity and curvature) (Chesarino et al., 2017; Guo et al., 2021; Rahman et al., 2020). Therefore, our results raise the likely scenario that cholesterol engagement by the AH is functionally tied to its ability to alter membranes in a manner that disfavors fusion between virus and cell. The link between cholesterol binding by the AH and its known functions is supported by the fact that F63Q and F67Q mutations, which were previously shown to cause loss of function of IFITM3 in cells and in artificial membranes (Chesarino et al., 2017; Guo et al., 2021), dramatically reduce cholesterol binding.

The AH is conserved among human IFITM family members, but distinct antiviral specificities have been demonstrated for IFITM1, IFITM2, and IFITM3. This is due, at least in part, to their differential subcellular localization. For example, IFITM1 is primarily located at the plasma membrane while IFITM2 and IFITM3 accumulate in endosomal membranes following endocytosis from the cell surface (Shi et al., 2017). We predict that IFITM1 and IFITM2 also engage cholesterol at their respective locations, and this may contribute to the block of virus entry at those sites. However, this prediction must be tested, for example by assessing whether the AH of IFITM1 and IFITM2 directly bind NBD-cholesterol *in vitro*. Furthermore, the trafficking of IFITM proteins is dynamic and controlled by post-translational modifications, including phosphorylation and S-palmitoylation (Chesarino et al., 2014b). The phosphorylation of IFITM3 negatively regulates its internalization from the plasma membrane by preventing endocytosis (Chesarino et al., 2014a; Compton et al., 2016; Jia et al., 2012). Furthermore, mutations in IFITM3 that prevent endocytosis similarly result in enhanced plasma membrane localization. IFITM3 at the cell surface performs important roles ranging from inhibition of HIV-1 infection (Compton et al., 2014; Compton et al., 2016; Lu et al., 2011) to promotion of PI3K signaling (Lee et al., 2020). Interestingly, depletion of IFITM3 from cells disrupts the formation of cholesterol-rich lipid rafts (Lee et al., 2020), which are important for both of these processes. Moreover, IFITM3 at the cell surface has been shown to promote infection of SARS-CoV-2, a virus that depends on cholesterol for entry into cells (Shi et al., 2020). Therefore, it will be interesting to ascertain how cholesterol binding contributes to the various functions, antiviral or otherwise, ascribed to IFITM3 and related IFITM proteins. The employment of additional methodologies, such as membrane floatation assays involving liposomes and recombinant IFITM3 protein, would be helpful in this regard.

In addition to a role played by cholesterol in the effector functions of IFITM3, the lipid may indirectly influence function by affecting the conformation and stability of IFITM3 in membranes. S-palmitoylation (the covalent addition of palmitic acid, a fatty acid) at conserved cysteines in IFITM proteins has been shown to facilitate membrane anchoring and extend protein half-life (Yount et al., 2012; Yount et al., 2010), but how cholesterol may affect IFITM3 localization and/or stability is unknown. It is unlikely that cholesterol binding itself impacts the degree to which IFITM3 is palmitoylated in cells, because it was previously shown that IFITM3 F67Q exhibits a similar degree of S-palmitoylation compared to WT (Chesarino et al., 2017). On the other hand, in the aforementioned pre-print posted during preparation of this manuscript, it was reported that a palmitoylation-deficient version of IFITM3 exhibited less cross-linking with a cholesterol analog in cells (Das et al., 2021). Therefore, the relationship between S-palmitoylation and cholesterol binding remains unclear and should be a focus of further mechanistic studies of IFITM function.

While we do not know how cholesterol binding by the AH of IFITM3 may regulate its functions, work performed on similar helices identified in other proteins provide important clues and future directions. For example, the M2 protein of IAV encodes an AH that also exhibits cholesterol binding activity (Ekanayake et al., 2016; Elkins et al., 2017; Martyna et al., 2020). In fact, the aromatic rings of membrane-facing phenylalanines within the AH are believed to play an important role in the M2-cholesterol interaction (Ekanayake et al., 2016). As a result, cholesterol binding by the AH of M2 increases its depth in lipid bilayers as well as its orientation and promotes membrane rigidity and curvature necessary for virus budding from the plasma membrane. Therefore, we suspect that engagement of cholesterol by the AH of IFITM3 similarly increases its penetrative depth and positioning within membranes and confers it with a greater capacity to stiffen and bend membranes during the virus-cell membrane fusion reaction. Indeed, our molecular dynamics simulations suggest that mutations that disrupt AH formation and cholesterol binding potential inhibit peptide embedding into membrane leaflets. Furthermore, in the aforementioned pre-print, the authors used molecular simulations to suggest that cholesterol affects the positioning of the AH of IFITM3 in membranes (Das et al., 2021). Therefore, our description of the cholesterol binding potential of the AH likely contributes to these atomic-level observations.

While we present clear evidence that the AH of IFITM3 is capable of binding cholesterol in a manner that requires phenylalanines, this region does not encode a clear cholesterol recognition motif (CRAC or CARC). However, this is not surprising since cholesterol binding has been demonstrated by many proteins lacking these motifs, and, importantly, the presence of these motifs does not necessarily predict cholesterol binding potential (Fantini and Barrantes, 2013; Taghon et al., 2021). Rather, the AH of IFITM3 may correspond to a “tilted” peptide that achieves functional lipid interactions due to its angular insertion in the membrane and the presence of membrane-facing phenylalanines. On the other hand, IFITM3 contains at least one cholesterol recognition motif (^104^<u>K</u>CLNI<u>W</u>ALI<u>L</u>^113^), and we show here that this site enables some degree of cholesterol binding *in vitro*. However, since the AH is necessary to modify membranes in living cells (Rahman et al., 2020) and sufficient to modify artificial membranes *in vitro* (Guo et al., 2021), the functional consequences of cholesterol binding elsewhere in IFITM3 are unclear. Perhaps the interaction between cholesterol and ^104^<u>K</u>CLNI<u>W</u>ALI<u>L</u>^113^ affects the depth and orientation of the AH in the context of the full-length IFITM3 protein, and there is some evidence to support this possibility (Das et al., 2021). Nonetheless, ^104^<u>K</u>CLNI<u>W</u>ALI<u>L</u>^113^ is not conserved in mouse IFITM3, despite being well characterized as a restriction factor against IAV in this species (Bailey et al., 2012; Everitt et al., 2012). In contrast, the AH of IFITM3 is more highly conserved across vertebrates and is known to alter the biomechanical properties of membranes on its own. Therefore, cholesterol binding by the AH of IFITM3 may represent an important feature contributing to its evolutionary conservation.

Furthermore, our work also has implications for the controversial role of VAPA as a co-factor for IFITM3 function (Amini-Bavil-Olyaee et al., 2013). Since the AH and the crucial phenylalanines at residues 63 and 67 found therein do not overlap with the region of IFITM3 that mediates an interaction with VAPA (the TMD) (Amini-Bavil-Olyaee et al., 2013), it is unlikely that mutations within the AH influence VAPA binding. While our results do not rule out that VAPA is involved in the functions of IFITM3, they indicate that accumulation of cholesterol in late endosomes per se is unlikely to be responsible for the antiviral activity of IFITM3. Instead, they suggest that the coincidence of cholesterol and IFITM3 at membrane sites serving as portals for virus entry is responsible for restriction. A model whereby IFITM3 and cholesterol are both required to inhibit fusion pore formation reconciles previously conflicting data from multiple publications. Specifically, it has been shown that enforced cholesterol accumulation in late endosomes following NPC1 inactivation variably inhibits IAV, which undergoes pH-dependent fusion at this site (Desai et al., 2014; Lin et al., 2013; Wrensch et al., 2014). In contrast, it is possible that loss of NPC1 function in cells expressing IFITM3 may result in virus entry inhibition. This possibility was supported by showing that inactivation of NPC1 by U18666A inhibits IAV infection in IFITM3-competent cells but lesser so in IFITM3-deficient cells (Kühnl et al., 2018). Nonetheless, direct evidence that IFITM3 function requires the presence of cholesterol demands methods that remove or inactivate cholesterol from cells. The use of methyl-beta-cyclodextrin to deplete cholesterol in IFITM3-expressing cells has been reported to have variable effects on infection (Amini-Bavil-Olyaee et al., 2013; Lin et al., 2013). Since many viruses depend upon cholesterol for entry (including IAV) (Sun and Whittaker, 2003), other strategies are needed in order to directly test a cholesterol requirement in the antiviral activities of IFITM3. In addition to affecting the angular depth of the AH in membranes, which may allow it to inflict change to membrane fluidity and curvature, it is possible that IFITM3 binds to and sequesters cholesterol that is ordinarily used by viral envelope glycoproteins to complete the membrane fusion reaction.

Overall, the findings presented and discussed here provide an advance towards our understanding of how IFITM3 impacts membrane microenvironments. However, it remains to be determined how coordination of cholesterol contributes to the precise mechanism by which IFITM3 disfavors fusion pore formation. A coordinated effort of *in silico, in vitro*, and *in cellulo* work will be required to resolve this important question and doing so will enable the development of innovative antiviral therapies that mimic the molecular action of IFITM3.

## Materials and Methods

### Peptide synthesis and reconstitution

Peptides listed in Table 1 were synthesized with >98% purity by Vivitide. Lyophilized peptides were reconstituted in DMSO or 30% acetonitrile containing 0.1% TFA to produce working stocks of 1 mg/mL and were stored at -20°C.

### Analysis of peptide binding to NBD-modified lipid

22-(N-(7-Nitrobenz-2-Oxa-1,3-Diazol-4-yl)Amino)-23,24-Bisnor-5-Cholen-3β-Ol-cholesterol (22-NBD-cholesterol) (N1148) and 7-NBD-PE (N360) were obtained from Thermo Fisher and dissolved in ethanol at 1 mM. Binding of peptides to NBD-cholesterol/NBD-PE was measured by either combining increasing concentrations of peptides (3.125 to 50 μM or 0.6125 to 5 μM) with 50-500 nM of NBD-cholesterol/NBD-PE or by combining increasing concentrations of NBD-cholesterol (0.00038 to 25 μM) with 5 μM of peptides. Reactions were carried out in 96-well black flat-bottom plates for 16 hours (unless indicated otherwise) at 4L in the presence of 20 mM HEPES (pH 7.4), 150 mM NaCl, 0.004% NP-40, 2% ethanol (v/v) and 2mM TCEP.

NBD fluorescence intensities were measured at 22°C within 5 mins of removal from 4L by fluorescence spectroscopy (excitation: 470 nm; emission: 540 nm) (Reitz et al., 2008) using a Tecan Infinite M1000. Fluorescence intensities obtained from NBD-cholesterol/NBD-PE alone (no peptide) were subtracted from fluorescence obtained from samples containing both NBD-cholesterol/NBD-PE and peptide. To derive the dissociation constant (*K*_*d*_) for NBD-cholesterol binding to P2, fluorescence measurements were fitted to a nonlinear regression curve in Prism. The least squares regression method was used under the assumption that total binding to a single site is the sum of specific and non-specific binding. Background was constrained to zero. For competition assays, 50 μM of unmodified cholesterol (C8667; Sigma) was pre-incubated with peptides prior to addition of 500 nM NBD-cholesterol, and one hour later, NBD fluorescence intensity was measured. NBD fluorescence polarization was measured in reactions as outlined above, except reactions were incubated at room temperature for 1 h prior to excitation with plane-polarized light using a Tecan Infinite M1000 (excitation: 470 nm; emission: 540 nm).

### Analysis of recombinant protein binding to NBD-cholesterol

Recombinant Glutathione-S-Transferase (GST) was obtained from Rockland Immunochemicals (001-001-200) and GST-IFITM3 was obtained from Abnova (H00010410-P01). GST was received at 1 mg/mL concentration in 0.2 M potassium phosphate, 0.15 M NaCl (pH 7.2) and GST-IFITM3 was received at 0.14 mg/mL concentration in 50 mM Tris-HCl and 10 mM glutathione. Binding of proteins to NBD-cholesterol was measured by combining increasing concentrations of protein (0.11 to 1.75 μM) with 500 nM NBD-cholesterol. Reactions were carried out in 96-well black flat-bottom plates for 1 hour at 4L in the presence of 20 mM HEPES (pH 7.4), 150 mM NaCl, 0.004% NP-40, and 2% ethanol (v/v). NBD fluorescence intensities were measured at 22°C within 5 mins of removal from 4L by fluorescence spectroscopy (excitation: 470 nm; emission: 540 nm) using a Tecan Infinite M1000. Fluorescence intensities obtained from NBD-cholesterol (no protein) were subtracted from fluorescence obtained from samples containing both NBD-cholesterol and protein.

### Western blot analysis

2 µg of GST or GST-IFITM3 protein was loaded into a 12% acrylamide Criterion XT Bis-Tris Precast Gel (Bio-Rad). Electrophoresis was performed with NuPage MES SDS Running Buffer (Invitrogen) and proteins were transferred to Amersham Protran Premium Nitrocellulose Membrane, pore size 0.20 µm (GE Healthcare). Membranes were blocked with Odyssey Blocking Buffer (Li-COR) and incubated with the following primary antibodies diluted in Odyssey Antibody Diluent (Li-COR): anti-GST (sc-138; Santa Cruz Biotechnology) and anti-IFITM3 (EPR5242, ab109429; Abcam). Secondary antibodies conjugated to DyLight 800 or 680 (Li-Cor) and the Li-Cor Odyssey CLx imaging system were used to reveal specific protein detection. Images were analyzed and assembled using ImageStudioLite (Li-Cor).

### Structural characterization of peptides using circular dichroism (CD)

25 μg of peptide was lyophilized and resuspended in 10 mM sodium borate (pH 7.4), 150 mM NaCl, 3.3% ethanol and 25 mM SDS to achieve a final peptide concentration of 60 μM and to promote a hydrophobic environment and peptide folding according to a previous report (Chesarino et al., 2017). A solution lacking peptide was used for background correction. Spectra were acquired at 25LC in continuous mode between 200 and 250 nm on a Jasco J-1500 CD Spectropolarimeter. The spectra were recorded at a scan rate of 10 nm/min, with a data pitch of 1 nm, a bandwidth of 1 nm, and a digital integration time of 8 seconds, and were presented as averages of three acquisitions. The spectra were deconvolved to estimate the secondary structural content using PEPFIT which uses a peptide-specific basis set (Reed and Reed, 1997). Averages of the top five deconvolved estimates for each peptide are presented.

### Intrinsic tryptophan fluorescence and FRET

50 μM P2 peptide was incubated in the presence or absence of NBD-cholesterol/NBD-PE for 16 hours at 4°C and intrinsic tryptophan fluorescence was measured by fluorescence spectroscopy (excitation: 295 nm; emission: 205-300 nm) (Reitz et al., 2008) using a Tecan Infinite M1000. FRET between peptide and NBD-cholesterol/NBD-PE was measured by fluorescence spectroscopy (excitation: 295 nm; emission: 500-600 nm) using a Tecan Infinite M1000.

### Transfection and virus infections

Influenza A Virus [A/PR/8/34 (PR8), H1N1] was purchased from Charles River Laboratories. HEK293T cells (ATCC: CRL-3216) were seeded in 12-well plates and transiently transfected with plasmid amounts that resulted in approximately equivalent mean fluorescence intensities of FLAG, as determined by flow cytometry (**Supplemental Figure 1A**): 0.75 μg pQCXIP-FLAG-PRRT2, 0.75 μg pQCXIP-FLAG-IFITM3 WT or 2.0 μg pQCXIP-FLAG-IFITM3-F63Q/F67Q/Δ59-68 using Lipofectamine 2000 (Invitrogen). At 24 h post-transfection, cells were re-plated in 24-well plates (50,000 per well). The following day, cells were overlaid with virus (MOI = 0.2) and at 18 h post-infection, cells were fixed and permeabilized with Cytofix/Cytoperm (BD), immunostained with anti-IAV NP (Abcam, AA5H), and analyzed on a LSRFortessa flow cytometer (BD).

### Molecular dynamics system build and simulations

WT and mutant peptides were built as ideal alpha-helices. For each peptide species, one peptide per leaflet was placed near the membrane-water interface (Jo et al., 2007; Lee et al., 2016). The membrane was composed of 70:30 mol% POPC:cholesterol with 200 lipids per leaflet, and the solution was 50 H_2_O per lipid with 150 mM KCl. Each system type was built in triplicate. The systems were minimized and briefly equilibrated using NAMD (Phillips et al., 2020) and then converted to Amber format (Case et al., 2005) using ParmEd. The simulations used the CHARMM36m all-atom force field (Huang et al., 2017; Klauda et al., 2010) and Amber20 version of pmemd.cuda (Case et al., 2020). See the methods described for previous planar membrane simulations for more detailed information (Beaven et al., 2022; Lessen et al., 2022). Briefly, constant pressure of 1 bar was maintained by a Monte Carlo barostat, and constant temperature was maintained at 310.15 K using Langevin dynamics with a 1 ps^−1^ damping coefficient. Non-bonded forces were switched off between 10-12 Å. Covalent bonds involving hydrogen were constrained using the SHAKE and SETTLE algorithms. Long-range electrostatics were calculated by particle mesh Ewald summation. All independent replicas were simulated for 1 µs, and the analysis used the last 0.9 µs of each run.

### Sequence retrieval and alignment

*IFITM3* sequences from the indicated species (common name shown) were retrieved from NCBI GenBank, partial amino acid alignments were generated using MUSCLE.

## Acknowledgements

Work in the lab of AAC is funded by the NIH Intramural Research Program, Center for Cancer Research, National Cancer Institute. This work utilized the computational resources of the NIH HPC Biowulf cluster (http://hpc.nih.gov). AHB was supported by a Postdoctoral Research Associate fellowship from the National Institute of General Medical Sciences, award number 1Fi2GM137844-01.

---

**Supplemental Figure 1:** NBD-cholesterol (500 nM) fluorescence intensities measured by spectroscopy (excitation: 470 nm; emission: 540 nm) following incubation with increasing concentrations (3.125 to 50 μM) of the indicated peptides. Incubation times prior to fluorescence reading were 1 hour, 2 hours, or 16 hours. Results represent the mean of three independent experiments and are normalized to 50 μM P2 peptide + NBD-cholesterol (set to 100%) for each timepoint. Error bars indicate standard error.

**Supplemental Figure 2:** (A) HEK293T cells transfected with PRRT2, IFITM3 WT, or IFITM3 mutants (F63Q or F67Q or Δ59-68) were fixed/permeabilized, stained with anti-FLAG, and the mean fluorescence intensity (MFI) of cells was measured by flow cytometry. Relative MFI is presented as a percentage of the MFI exhibited by IFITM3 WT. Results represent the mean of three independent experiments and are normalized to IFITM3 WT (set to 100%). Error bars indicate standard error. Inset shows representative overlaying dot plots from which MFI was calculated. (B) HEK293T cells were transfected with PRRT2, IFITM3 WT, or IFITM3 mutants (F63Q or F67Q or Δ59-68) and challenged with IAV (MOI = 0.2). At 18 h post-infection, cells were fixed and permeabilized, stained with anti-NP, and analyzed by flow cytometry. (C) NBD-cholesterol (50 nM) fluorescence polarization was measured by spectroscopy (excitation with plane-polarized light: 470 nm; emission: 540 nm) following incubation with increasing concentrations (0.625 to 5 μM) of the indicated peptides. Results represent the mean of three independent experiments and are normalized to 5 μM P2 peptide + NBD-cholesterol (set to 100%). (D) NBD-cholesterol (500 nM) fluorescence intensities measured by spectroscopy (excitation: 470 nm; emission: 540 nm) following 1 hour incubation with increasing concentrations (3.125 to 50 μM) of P2’ peptide. In the condition indicated, P2’ was pre-incubated with unmodified cholesterol (50 μM) for 3 hours followed by incubation with NBD-cholesterol (500 nM) for 1 hour. Results represent the mean of three independent experiments and are normalized to 50 μM P2’ + NBD-cholesterol (set to 100%). Error bars indicate standard error. Asterisks indicate statistically significant difference (p<0.05) between P2 and another peptide (condition indicated by asterisk color), as determined by one-way ANOVA (B & C) or student T-test (D).

## References

Amini-Bavil-Olyaee, S., Y.J. Choi, J.H. Lee, M. Shi, I.-C. Huang, M. Farzan, and J.U. Jung. 2013. The Antiviral Effector IFITM3 Disrupts Intracellular Cholesterol Homeostasis to Block Viral Entry. CHOM 13:452–464.

Avdulov, N.A., S.V. Chochina, U. Igbavboa, C.S. Warden, A.V. Vassiliev, and W.G. Wood. 1997. Lipid binding to amyloid beta-peptide aggregates: preferential binding of cholesterol as compared with phosphatidylcholine and fatty acids. J Neurochem 69:1746–1752.

Bailey, C.C., I.-C. Huang, C. Kam, and M. Farzan. 2012. Ifitm3 limits the severity of acute influenza in mice. PLoS Pathogens 8:e1002909.

Beaven, A.H., K. Sapp, and A.J. Sodt. 2022. Dynamic cholesterol redistribution favors membrane fusion pore constriction. bioRxiv

Carette, J.E., M. Raaben, A.C. Wong, A.S. Herbert, G. Obernosterer, N. Mulherkar, A.I. Kuehne, P.J. Kranzusch, A.M. Griffin, G. Ruthel, P. Dal Cin, J.M. Dye, S.P. Whelan, K. Chandran, and T.R. Brummelkamp. 2011. Ebola virus entry requires the cholesterol transporter Niemann-Pick C1. Nature 477:340–343.

Case, D.A., I.Y. Ben-Shalom, S.R. Brozell, D.S. Cerutti, T.E. Cheatham, I. Cruzeiro, V.W.D. T.A. Darden, R.E. Duke, D. Ghoreishi, M.K. Gilson, H. Gohlke, A.W. Goetz, D. Greene, R. Harris, N. Homeyer, S. Izadi, A. Kovalenko, T. Kurtzman, T.S. Lee, S. LeGrand, P. Li, C. Lin, J. Liu, T. Luchko, R. Luo, D.J. Mermelstein, K.M. Merz, Y. Miao, G. Monard, C. Nguyen, H. Nguyen, I. Omelyan, A. Onufriev, F. Pan, R. Qi, D.R. Roe, A. Roitberg, C. Sagui, S. Schott-Verdugo, J. Shen, C.L. Simmerling, J. Smith, R. Salomon-Ferrer, J. Swails, R.C. Walker, J. Wang, H. Wei, R.M. Wolf, X. Wu, L. Xiao, D.M. York, and P.A. Kollman. 2020. AMBER 2020, University of California.

Case, D.A., T.E. Cheatham, 3rd, T. Darden, H. Gohlke, R. Luo, K.M. Merz, Jr., A. Onufriev, C. Simmerling, B. Wang, and R.J. Woods. 2005. The Amber biomolecular simulation programs. J Comput Chem 26:1668–1688.

Chernomordik, L.V., and M.M. Kozlov. 2003. Protein-Lipid Interplay in Fusion and Fission of Biological Membranes. Annual Review of Biochemistry 72:175–207.

Chesarino, N.M., A.A. Compton, T.M. McMichael, A.D. Kenney, L. Zhang, V. Soewarna, M. Davis, O. Schwartz, and J.S. Yount. 2017. IFITM3 requires an amphipathic helix for antiviral activity. EMBO reports 18:e201744100–201744112.

Chesarino, N.M., T.M. McMichael, J.C. Hach, and J.S. Yount. 2014a. Phosphorylation of the antiviral protein interferon-inducible transmembrane protein 3 (IFITM3) dually regulates its endocytosis and ubiquitination. Journal of Biological Chemistry 289:11986–11992.

Chesarino, N.M., T.M. McMichael, and J.S. Yount. 2014b. Regulation of the trafficking and antiviral activity of IFITM3 by post-translational modifications. Future Microbiology 9:1151–1163.

Compton, A.A., T. Bruel, F. Porrot, A. Mallet, M. Sachse, M. Euvrard, C. Liang, N. Casartelli, and O. Schwartz. 2014. IFITM Proteins Incorporated into HIV-1 Virions Impair Viral Fusion and Spread. Cell Host & Microbe 16:736–747.

Compton, A.A., N. Roy, F. Porrot, A. Billet, N. Casartelli, J.S. Yount, C. Liang, and O. Schwartz. 2016. Natural mutations in IFITM3 modulate post-translational regulation and toggle antiviral specificity. EMBO reports 17:1657–1671.

Coomer, C.A., K. Rahman, and A.A. Compton. 2021. CD225 Proteins: A Family Portrait of Fusion Regulators. Trends in Genetics 7:1–4.

Das, T., X. Yang, H. Lee, E. Garst, E. Valencia, K. Chandran, W. Im, and H. Hang. 2021. S-palmitoylation and sterol interactions mediate antiviral specificity of IFITM isoforms. Res Sq

Desai, T.M., M. Marin, C.R. Chin, G. Savidis, A.L. Brass, and G.B. Melikyan. 2014. IFITM3 restricts influenza A virus entry by blocking the formation of fusion pores following virus-endosome hemifusion. PLoS Pathogens 10:e1004048.

Drin, G., and B. Antonny. 2010. Amphipathic helices and membrane curvature. FEBS Letters 584:1840–1847.

Ekanayake, E.V., R. Fu, and T.A. Cross. 2016. Structural Influences: Cholesterol, Drug, and Proton Binding to Full-Length Influenza A M2 Protein. Biophys J 110:1391–1399.

Elkins, M.R., J.K. Williams, M.D. Gelenter, P. Dai, B. Kwon, I.V. Sergeyev, B.L. Pentelute, and M. Hong. 2017. Cholesterol-binding site of the influenza M2 protein in lipid bilayers from solid-state NMR. Proc Natl Acad Sci U S A 114:12946–12951.

Everitt, A.R., S. Clare, T. Pertel, S.P. John, R.S. Wash, S.E. Smith, C.R. Chin, E.M. Feeley, J.S. Sims, D.J. Adams, H.M. Wise, L. Kane, D. Goulding, P. Digard, V. Anttila, J.K. Baillie, T.S. Walsh, D.A. Hume, A. Palotie, Y. Xue, V. Colonna, C. Tyler-Smith, J. Dunning, S.B. Gordon, K. Everingham, H. Dawson, D. Hope, P. Ramsay, T.S. Walsh Local Lead Investigator, A. Campbell, S. Kerr, D. Harrison, K. Rowan, J. Addison, N. Donald, S. Galt, D. Noble, J. Taylor, N. Webster Local Lead Investigator, I. Taylor Local Lead Investigator, J. Aldridge Local Lead Investigator, R. Dornan, C. Richard, D. Gilmour, R. Simmons Local Lead Investigator, R. White Local Lead Investigator, C. Jardine, D. Williams Local Lead Investigator, M. Booth Local Lead Investigator, T. Quasim, V. Watson, P. Henry, F. Munro, L. Bell, J. Ruddy Local Lead Investigator, S. Cole Local Lead Investigator, J. Southward, P. Allcoat, S. Gray, M. McDougall Local Lead Investigator, J. Matheson, J. Whiteside Local Lead Investigator, D. Alcorn, K. Rooney Local Lead Investigator, R. Sundaram, G. Imrie Local Lead Investigator, J. Bruce, K. McGuigan, S. Moultrie Local Lead Investigator, C. Cairns Local Lead Investigator, J. Grant, M. Hughes, C. Murdoch Local Lead Investigator, A. Davidson Local Lead Investigator, G. Harris, R. Paterson, C. Wallis Local Lead Investigator, S. Binning Local Lead Investigator, M. Pollock, J. Antonelli, A. Duncan, J. Gibson, C. McCulloch, L. Murphy, C. Haley, G. Faulkner, T. Freeman, D.A. Hume, J.K. Baillie Principal Investigator, D. Chaussabel, W.E. Adamson, W.F. Carman, C. Thompson, M.C. Zambon, P. Aylin, D. Ashby, W.S. Barclay, S.J. Brett, W.O. Cookson, L.N. Drumright, J. Dunning, R.A. Elderfield, L. Garcia-Alvarez, B.G. Gazzard, M.J. Griffiths, M.S. Habibi, T.T. Hansel, J.A. Herberg, A.H. Holmes, T. Hussell, S.L. Johnston, O.M. Kon, M. Levin, M.F. Moffatt, S. Nadel, P.J. Openshaw, J.O. Warner, S.J. Aston, S.B. Gordon, A. Hay, J. McCauley, A. O’Garra, J. Banchereau, A. Hayward, P. Kellam, J.K. Baillie, D.A. Hume, P. Simmonds, P.S. McNamara, M.G. Semple, R.L. Smyth, J.S. Nguyen-Van-Tam, L.P. Ho, A.J. McMichael, R.L. Smyth, P.J. Openshaw, G. Dougan, A.L. Brass, and P. Kellam. 2012. IFITM3 restricts the morbidity and mortality associated with influenza. Nature 484:519–523.

Fantini, J., and F.J. Barrantes. 2013. How cholesterol interacts with membrane proteins: an exploration of cholesterol-binding sites including CRAC, CARC, and tilted domains. Front Physiol 4:31.

Guo, X., J. Steinkühler, M. Marin, X. Li, W. Lu, R. Dimova, and G.B. Melikyan. 2021. Interferon-Induced Transmembrane Protein 3 Blocks Fusion of Diverse Enveloped Viruses by Altering Mechanical Properties of Cell Membranes. ACS nano

Huang, J., S. Rauscher, G. Nawrocki, T. Ran, M. Feig, B.L. de Groot, H. Grubmuller, and A.D. MacKerell, Jr. 2017. CHARMM36m: an improved force field for folded and intrinsically disordered proteins. Nat Methods 14:71–73.

Jia, R., Q. Pan, S. Ding, L. Rong, S.-L. Liu, Y. Geng, W. Qiao, and C. Liang. 2012. The N-terminal region of IFITM3 modulates its antiviral activity by regulating IFITM3 cellular localization. Journal of Virology 86:13697–13707.

Jo, S., T. Kim, and W. Im. 2007. Automated builder and database of protein/membrane complexes for molecular dynamics simulations. PLoS One 2:e880.

Klauda, J.B., R.M. Venable, J.A. Freites, J.W. O’Connor, D.J. Tobias, C. Mondragon-Ramirez, I. Vorobyov, A.D. MacKerell, Jr., and R.W. Pastor. 2010. Update of the CHARMM all-atom additive force field for lipids: validation on six lipid types. J Phys Chem B 114:7830–7843.

Kühnl, A., A. Musiol, N. Heitzig, D.E. Johnson, C. Ehrhardt, T. Grewal, V. Gerke, S. Ludwig, and U. Rescher. 2018. Late Endosomal/Lysosomal Cholesterol Accumulation Is a Host Cell-Protective Mechanism Inhibiting Endosomal Escape of Influenza A Virus. mBio 9:665–616.

Lee, J., X. Cheng, J.M. Swails, M.S. Yeom, P.K. Eastman, J.A. Lemkul, S. Wei, J. Buckner, J.C. Jeong, Y. Qi, S. Jo, V.S. Pande, D.A. Case, C.L. Brooks, 3rd, A.D. MacKerell, Jr., J.B. Klauda, and W. Im. 2016. CHARMM-GUI Input Generator for NAMD, GROMACS, AMBER, OpenMM, and CHARMM/OpenMM Simulations Using the CHARMM36 Additive Force Field. J Chem Theory Comput 12:405–413.

Lee, J., M.E. Robinson, N. Ma, D. Artadji, M.A. Ahmed, G. Xiao, T. Sadras, G. Deb, J. Winchester, K.N. Cosgun, H. Geng, L.N. Chan, K. Kume, T.P. Miettinen, Y. Zhang, M.A. Nix, L. Klemm, C.W. Chen, J. Chen, V. Khairnar, A.P. Wiita, A. Thomas-Tikhonenko, M. Farzan, J.U. Jung, D.M. Weinstock, S.R. Manalis, M.S. Diamond, N. Vaidehi, and M.M.x.F. schen. 2020. IFITM3 functions as a PIP3 scaffold to amplify PI3K signalling in B cells. Nature 1:1–35.

Lessen, H.J., K.C. Sapp, A.H. Beaven, R. Ashkar, and A.J. Sodt. 2022. Molecular mechanisms of spontaneous curvature and softening in complex lipid bilayer mixtures. bioRxiv

Li, K., R.M. Markosyan, Y.-M. Zheng, O. Golfetto, B. Bungart, M. Li, S. Ding, Y. He, C. Liang, J.C. Lee, E. Gratton, F.S. Cohen, and S.-L. Liu. 2013. IFITM proteins restrict viral membrane hemifusion. PLoS Pathogens 9:e1003124.

Lin, T.-Y., C.R. Chin, A.R. Everitt, S. Clare, J.M. Perreira, G. Savidis, A.M. Aker, S.P. John, D. Sarlah, E.M. Carreira, S.J. Elledge, P. Kellam, and A.L. Brass. 2013. Amphotericin B Increases Influenza A Virus Infection by Preventing IFITM3-Mediated Restriction. Cell Reports 5:895–908.

Loura, L.M., A. Fedorov, and M. Prieto. 2001. Exclusion of a cholesterol analog from the cholesterol-rich phase in model membranes. Biochim Biophys Acta 1511:236–243.

Lu, J., Q. Pan, L. Rong, W. He, S.-L. Liu, and C. Liang. 2011. The IFITM proteins inhibit HIV-1 infection. Journal of Virology 85:2126–2137.

Majdoul, S., and A.A. Compton. 2021. Lessons in self-defence: inhibition of virus entry by intrinsic immunity. Nature Reviews Immunology

Martyna, A., B. Bahsoun, J.J. Madsen, F. Jackson, M.D. Badham, G.A. Voth, and J.S. Rossman. 2020. Cholesterol Alters the Orientation and Activity of the Influenza Virus M2 Amphipathic Helix in the Membrane. J Phys Chem B 124:6738–6747.

Omasits, U., C.H. Ahrens, S. Müller, and B. Wollscheid. 2014. Protter: interactive protein feature visualization and integration with experimental proteomic data. Bioinformatics 30:884–886.

Phillips, J.C., D.J. Hardy, J.D.C. Maia, J.E. Stone, J.V. Ribeiro, R.C. Bernardi, R. Buch, G. Fiorin, J. Henin, W. Jiang, R. McGreevy, M.C.R. Melo, B.K. Radak, R.D. Skeel, A. Singharoy, Y. Wang, B. Roux, A. Aksimentiev, Z. Luthey-Schulten, L.V. Kale, K. Schulten, C. Chipot, and E. Tajkhorshid. 2020. Scalable molecular dynamics on CPU and GPU architectures with NAMD. J Chem Phys 153:044130.

Poh, M.K., G. Shui, X. Xie, P.-Y. Shi, M.R. Wenk, and F. Gu. 2012. U18666A, an intra-cellular cholesterol transport inhibitor, inhibits dengue virus entry and replication. Antiviral Research 93:191–198.

Rahman, K., C.A. Coomer, S. Majdoul, S.Y. Ding, S. Padilla-Parra, and A.A. Compton. 2020. Homology-guided identification of a conserved motif linking the antiviral functions of IFITM3 to its oligomeric state. eLife 9:975–925.

Reed, J., and T.A. Reed. 1997. A set of constructed type spectra for the practical estimation of peptide secondary structure from circular dichroism. Anal Biochem 254:36–40.

Reitz, J., K. Gehrig-Burger, J.F. Strauss, 3rd, and G. Gimpl. 2008. Cholesterol interaction with the related steroidogenic acute regulatory lipid-transfer (START) domains of StAR (STARD1) and MLN64 (STARD3). FEBS J 275:1790–1802.

Schroeder, M.E., H.A. Hostetler, F. Schroeder, and J.M. Ball. 2012. Elucidation of the Rotavirus NSP4-Caveolin-1 and -Cholesterol Interactions Using Synthetic Peptides. J Amino Acids 2012:575180.

Shi, G., A.D. Kenney, E. Kudryashova, A. Zani, L. Zhang, K.K. Lai, L. Hall-Stoodley, R.T. Robinson, D.S. Kudryashov, A.A. Compton, and J.S. Yount. 2020. Opposing activities of IFITM proteins in SARSLCoVL2 infection. The EMBO journal 3:e201900542–201900512.

Shi, G., O. Schwartz, and A.A. Compton. 2017. More than meets the I: the diverse antiviral and cellular functions of interferon-induced transmembrane proteins. Retrovirology 14:1–11.

Sodt, A.J., and R.W. Pastor. 2014. Molecular modeling of lipid membrane curvature induction by a peptide: more than simply shape. Biophys J 106:1958–1969.

Sun, X., and G.R. Whittaker. 2003. Role for influenza virus envelope cholesterol in virus entry and infection. J Virol 77:12543–12551.

Taghon, G.J., J.B. Rowe, N.J. Kapolka, and D.G. Isom. 2021. Predictable cholesterol binding sites in GPCRs lack consensus motifs. Structure 29:499–506 e493.

Teissier, E., and E.-I. Pécheur. 2007. Lipids as modulators of membrane fusion mediated by viral fusion proteins. European Biophysics Journal 36:887–899.

Vivian, J.T., and P.R. Callis. 2001. Mechanisms of tryptophan fluorescence shifts in proteins. Biophys J 80:2093–2109.

Wrensch, F., M. Winkler, and S. Pöhlmann. 2014. IFITM Proteins Inhibit Entry Driven by the MERS-Coronavirus Spike Protein: Evidence for Cholesterol-Independent Mechanisms. Viruses 6:3683–3698.

Wustner, D. 2007. Fluorescent sterols as tools in membrane biophysics and cell biology. Chem Phys Lipids 146:1–25.

Yount, J.S., R.A. Karssemeijer, and H.C. Hang. 2012. S-palmitoylation and ubiquitination differentially regulate interferon-induced transmembrane protein 3 (IFITM3)-mediated resistance to influenza virus. Journal of Biological Chemistry 287:19631–19641.

Yount, J.S., B. Moltedo, Y.-Y. Yang, G. Charron, T.M. Moran, C.B. LÃ<sup>3</sup>pez, and H.C. Hang. 2010. Palmitoylome profiling reveals S-palmitoylation-dependent antiviral activity of IFITM3. Nature chemical biology 6:610–614.

Zhao, X., J. Li, C.A. Winkler, P. An, and J.T. Guo. 2018. IFITM Genes, Variants, and Their Roles in the Control and Pathogenesis of Viral Infections. Front Microbiol 9:3228.

