## Supplementary figures and images for "Cholesterol binds the amphipathic helix of IFITM3 and regulates antiviral activity"

### Supplemental Figures

Supplemental Figure 1

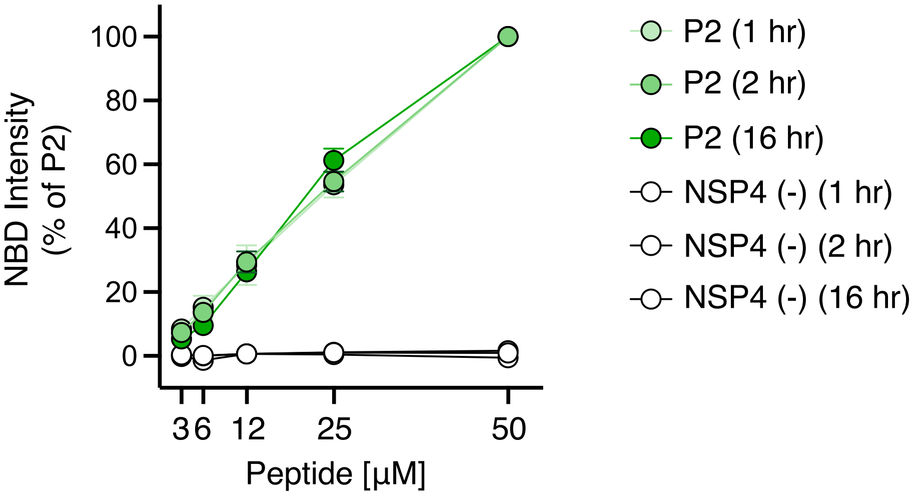

Supplemental Figure 2

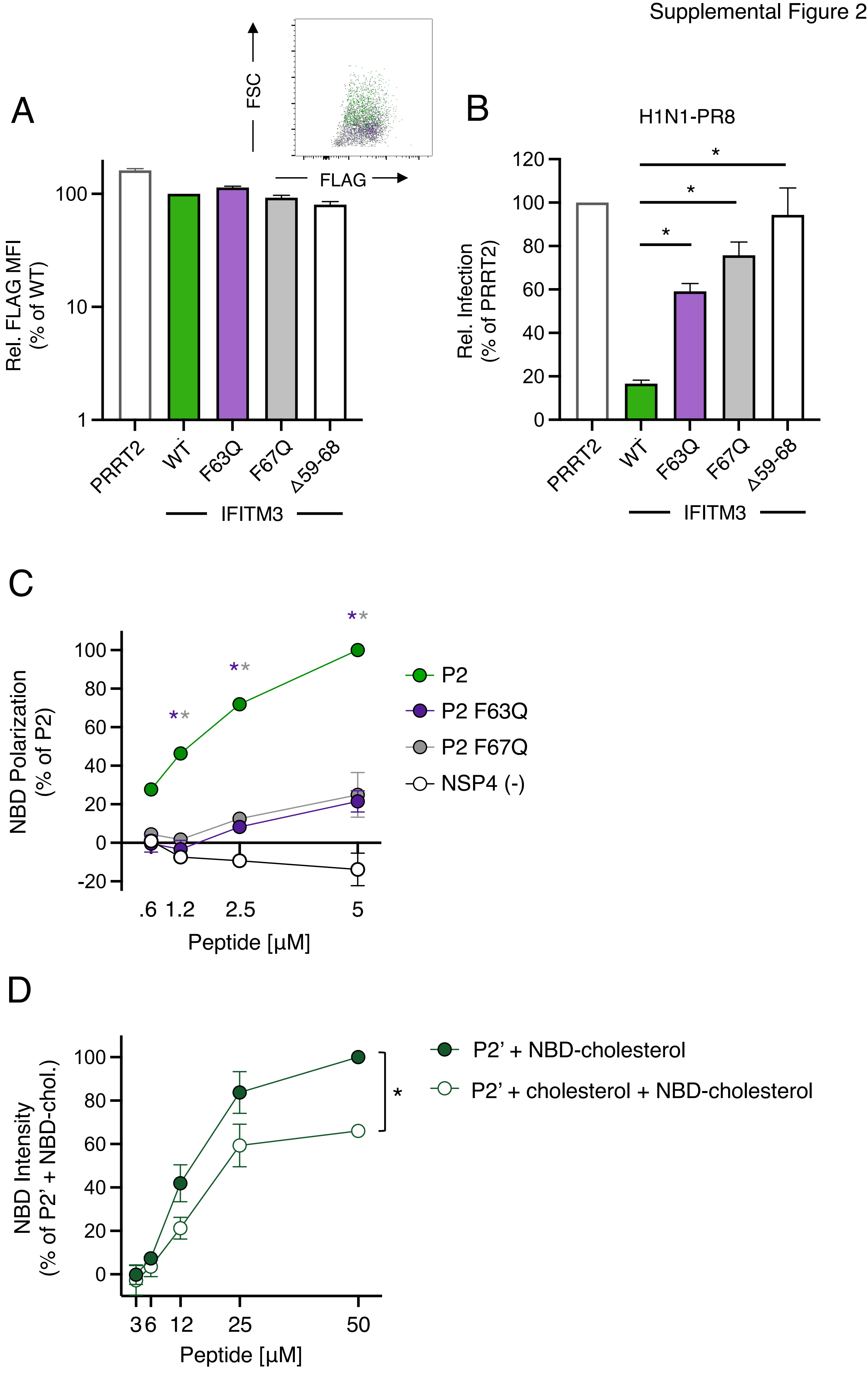
